# Antibody gene features associated with binding and functional activity in vaccine-derived human mAbs targeting malaria parasites

**DOI:** 10.1101/2023.08.01.551554

**Authors:** Camila H. Coelho, Susanna Marquez, Bergeline C. Nguemwo Tentokam, Anne D. Berhe, Kazutoyo Miura, Carole A. Long, Issaka Sagara, Sara Healy, Steven H. Kleinstein, Patrick E. Duffy

**Affiliations:** Laboratory of Malaria Immunology and Vaccinology, National Institute of Allergy and Infectious Diseases, National Institutes of Health, Bethesda, MD 20892, USA; Department of Microbiology, Icahn School of Medicine at Mount Sinai, New York, 10029, NY; Department of Pathology, Yale School of Medicine, New Haven, CT 06520, USA; Laboratory of Malaria and Vector and Research, National Institute of Allergy and Infectious Diseases, NIH, Rockville, Maryland, USA; Malaria Research and Training Center, University of Sciences, Techniques, and Technology, Bamako, Mali; Program in Computational Biology and Bioinformatics, Yale University, New Haven, CT 06511, USA; Department of Immunobiology, Yale School of Medicine, New Haven, CT 06520, USA

## Abstract

Adjuvants have been essential to malaria vaccine development, but their impact on the vaccine-induced antibody repertoire is poorly understood. Here, we used cDNA sequences from antigen-specific single memory B cells to express 132 recombinant human anti-Pfs230 monoclonal antibodies (mAbs). Alhydrogel®-induced mAbs demonstrated higher binding to Pfs230D1, although functional activity was similar between adjuvants. All Alhydrogel® mAbs using IGHV1-69 gene bound to recombinant Pfs230D1, but none blocked parasite transmission to mosquitoes; similarly, no AS01 mAb using IGHV1-69 blocked transmission. Functional mAbs from both Alhydrogel® and AS01 vaccines used IGHV3-21 and IGHV3-30 genes. Antibodies with the longest CDR3 sequences were associated with binding but not functional activity. This study assesses adjuvant effects on antibody clonotype diversity during malaria vaccination.

## INTRODUCTION

Despite intense efforts to eradicate malaria, the disease still affects hundreds of millions of people worldwide, of which over 90% live in Sub-Saharan Africa. Recent progress in malaria control has stalled, and in some areas, reversed,^1^ highlighting the need for new prevention tools. Although malaria vaccines are in advanced stages of clinical trials or even reaching implementation,^2,3^ long-term high-level protection has not yet been achieved. Many malaria vaccine candidates rely on antibody effector mechanisms and depend on newer adjuvants, thus the antibody repertoire response to individual *Plasmodium* proteins and newer adjuvants requires characterization to guide vaccine improvement.

Alhydrogel® is an adjuvant comprised of aluminum hydroxide gel, widely used in licensed vaccines for almost a century.^4^ Alhydrogel® generally induces a Th2-type immune response, and is often considered the “benchmark” adjuvant in comparative human studies with new adjuvants.^5^ However, novel adjuvants developed in recent decades have been included in licensed vaccines.^6,7^ AS01 is a novel adjuvant system that incorporates the TLR4 ligand, monophosphoryl lipid A (MPL), and the saponin fraction QS21 in a liposomal preparation.^8^ AS01 efficiently promotes humoral and cellular immune responses, including CD4 T cells, that in turn further assist B cells to generate high affinity antibodies. A recent study highlighted the ability of AS01 to enhance the antibody response versus Aluminum-based adjuvants. They measured increased antibody-associated signatures within whole blood transcriptomes using linear regression and linking 300 genes to antibody response.^9^

Previous studies have shown that Alhydrogel® might not be efficient when used in influenza vaccine formulations,^10,11^ whereas oil-in-water adjuvants, such as MF59 and AS03, induced higher antibody titers and neutralization rates against influenza viruses.^12,13^ Additionally, several malaria vaccine candidates failed clinical trials when formulated with Alhydrogel®-based adjuvants. Thus, improved vaccines are being developed and employ novel adjuvants, such as the recently approved pediatric RTS,S vaccine (Mosquirix, GSK) that uses AS01E adjuvant similar to the AS01B used in the adult shingles vaccine Shingrix from GSK.

Conventional serological approaches used to characterize immune response during malaria vaccination do not define properties of the antibody genes elicited in response to a given vaccine or adjuvant. Here, we sought to gain this insight by sorting antigen-specific memory B cells from healthy Malian adults immunized with Pfs230D1, a malaria transmission-blocking vaccine candidate targeting the parasite gamete stage, formulated with either Alhydrogel® or AS01.

## RESULTS

To assess the relationship between V gene usage and antibody binding or functional activity, we expressed 132 human IgG1 recombinant monoclonal antibodies (mAbs) using cDNA sequences of single Pfs230D1-specific memory B cells. We then tested their capacity for binding Pfs230D1 (by ELISA) and for functional activity (Figure **1a**). Function of antibodies was reported as TRA (*Transmission Reducing Activity)*, measured as the capacity of the mAb to reduce the parasite (oocyst) burden in mosquitoes a week after feeding on infective blood,^14,15^ an assay referred to as Standard Membrane Feeding Assay or SMFA. Functional mAbs were defined here as those that achieved TRA >75% at 100μg/mL in the infective bloodmeal. The binding epitopes of selected functional antibodies were recently reported.^16^ The panel comprised 35 mAbs derived from Alhydrogel® recipients and 97 from AS01 recipients. Sequences were selected for mAb expression based on evidence of clonal expansion, as previously reported.^14^ The binding rate of mAbs recognizing Pfs230D1 (domain 1 of Pfs230 protein) in ELISAs was substantially higher in the Alhydrogel® group (32/35, 91.4%) than in the AS01 group (33/97, 34.0%).

**Figure 1.**
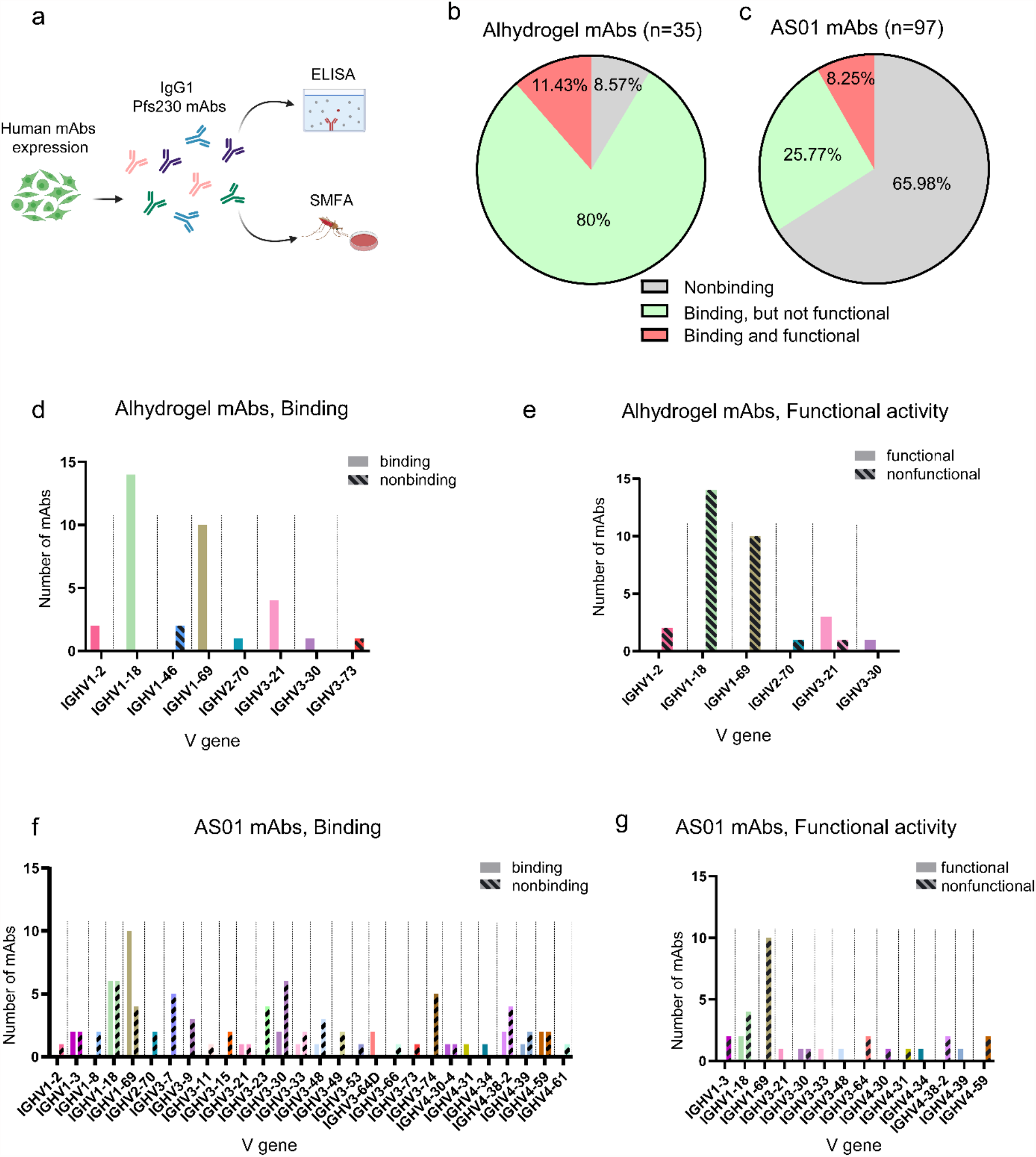
Binding and functional activity of Pfs230D1 mAbs and their relationship with heavy chain V gene usage. **(a)** Human IgG1 mAbs obtained from Pfs230D1-single B cells were expressed in mammalian cells and tested for binding to domain 1 of Pfs230 (ELISA). Functional activity was determined as Transmission Reducing Activity (TRA), assessed by Standard Membrane Feeding Assay using NF54 *Plasmodium falciparum* fed to *Anopheles* mosquitoes. Figure was created using Biorender.com **(b)** Percentage of human mAbs obtained from Alhydrogel® and **(c)** AS01 vaccinees that bound to recombinant Pfs230D1 and were functional in vivo, presenting TRA higher than 75% at 100μg/mL. **(d**,**e)** Alhydrogel and AS01 mAbs were grouped into binding and nonbinding and **(f**,**g)** functional and non-functional and then visualized according to their heavy chain V gene.

The overall percentages of functional antibodies were similar, 11.4% for Alhydrogel and 8.2% for AS01 (Figures **1b**,**c**). However, when analyzing the number of functional mAbs among Pfs230D1-binding antibodies, this number was higher for AS01 than for Alhydrogel®: 8 out of 33 (24.2%) AS01 mAbs were functional versus only 4 out of 32 (12.5%) Alhydrogel mAbs.

Interestingly, while all Alhydrogel® mAbs using IGHV1-18 (14/14) or IGHV1-69 (10/10) in their heavy chain bound successfully to recombinant Pfs230D1 (Figure **1d**), none of these were functional mAbs (Figure **1e**). The 97 mAbs from the AS01 group included 33 Pfs230D1-binding antibodies that were distributed among 14 different V genes in the IGHV1, IGHV3, and IGHV4 gene families (Figure **1f**). Within the AS01 recipients, IGHV1-18 and IGHV1-69 genes were most frequent, being used by 18% and 30% of binding antibodies, respectively (Figure **1f**). Among the 8 functional mAbs obtained from AS01 subjects, 2 were from IGHV1 gene family, 4 from IGHV3, and 2 from IGHV4 (Figure **1f**). Notably, both Alhydrogel® and AS01 vaccines generated functional mAbs using IGHV3-21 and IGHV3-30 genes (Figures **1e**,**g**).

Thirteen binding mAbs from the Alhydrogel group used IGHD6-13 genes in their sequence (Supplementary Figure **1**). Interestingly, 12/14 (85.7%) of the mAbs using IGHV1-18 also used IGHD6-13 genes (which represented 12/13 (92.3%) mAbs containing IGHD6-13 genes). Despite their ability to bind to recombinant antigen, none of the mAbs containing IGHD6-13 genes was functional (Supplementary Figure **1**). mAbs derived from AS01 recipients used diverse D genes and no specific gene appeared to be more frequent among the binding or functional antibodies (Supplementary Figure **1**).

For J genes used by Alhydrogel® mAbs, the majority of Pfs230D1-binding antibodies used IGHJ3 and IGHJ4 genes, while the few functional antibodies used IGHJ2 (2 mAbs) and IGHJ4 (2 mAbs) (Supplementary Figure **2)**. For AS01 mAbs, IGHJ4 was the most frequently used J gene family among Pfs230D1-binding and functional antibodies (19/33), followed by IGHJ5 and IGHJ6. (Supplementary Figure **2**).

In the Alhydrogel® response, CDR3 length (measured as the number of amino acids, aa) was higher for binding mAbs compared to nonbinding mAbs (p<0.005) (14 and 9 aa, respectively). The longest CDR3 sequence of Alhydrogel® mAbs was identified in a binding antibody (18 aa) while the shortest was in a nonbinding antibody (8 aa). For AS01 mAbs, binding and nonbinding had an average CDR3 length of 15 and 14 aa, respectively (p=0.09). For AS01 mAbs the longest CDR3 was found in binding mAbs (21 aa) and the shortest in nonbinding (7 aa).

No association was determined between functional activity and CDR3 length (Figure **2a**). Average CDR3 length for functional and nonfunctional mAbs was 13 aa and 14 aa respectively after Alhydrogel®, versus 15 aa and 15 aa after AS01 (Figure **2b**). Similar results were obtained after comparing functional vs. nonfunctional antibodies among only the mAbs known to bind antigen (Figure **2c**).

**Figure 2.**
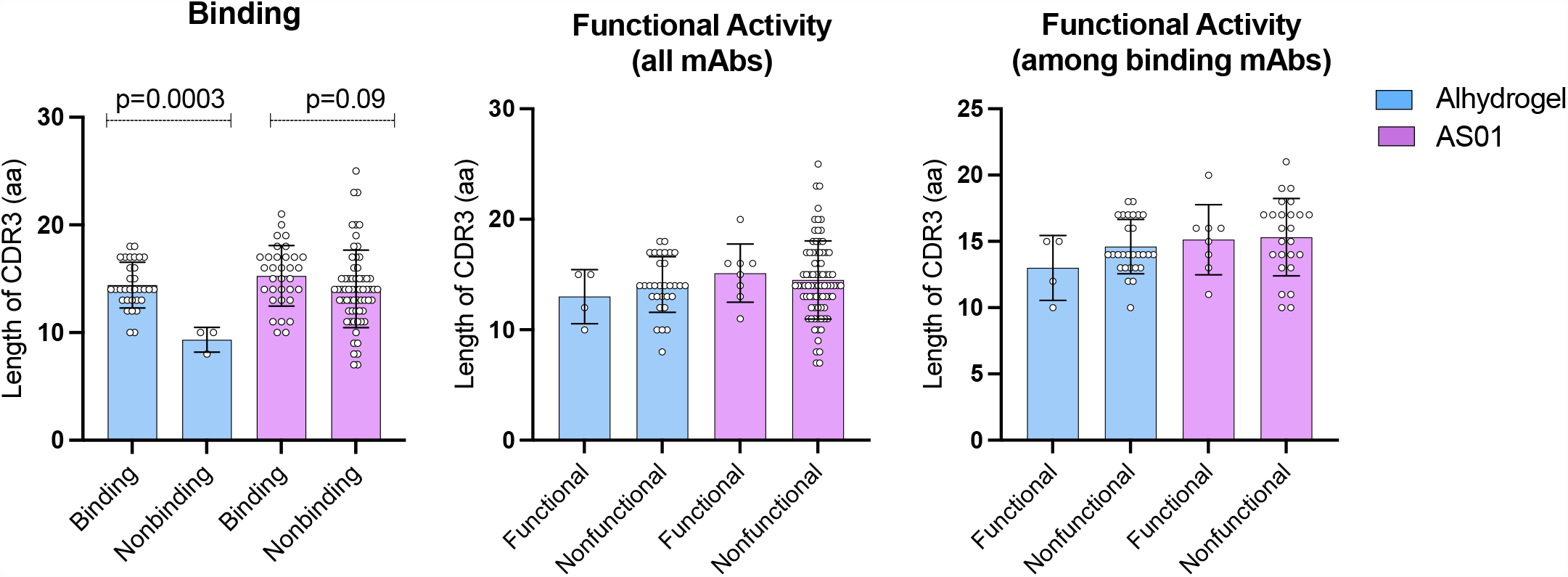
Association between heavy chain CDR3 length and binding or function of Pfs230D1 mAbs. **(a)** Binding was assessed by ELISA and compared between binding vs. non-binding mAbs in the two trials. **(b**,**c)** Functional activity was determined by Standard Membrane Feeding Assay (SMFA) and compared between functional and non-functional mAbs among **(b)** all mAbs, or **(c)** only those mAbs that bound antigen. P values were considered significant when less than 0.05 and statistical analyses was performed using unpaired t test. Mean and standard deviation are shown.

## DISCUSSION

Malaria TBVs aim to reduce parasitic burden in mosquitos carrying the pathogen *Plasmodium*, thus reducing malaria transmission.^17,18^ The leading TBV target antigen is Pfs230 (a 230 kDa protein present in the surface of *P. falciparum* gametes). Vaccine formulations containing domain 1 of Pfs230 adjuvanted with Alhydrogel® or AS01 are currently under evaluation in field trials in malaria-endemic areas (^19^, NCT05135273, NCT03917654), after a recent successful clinical trial in malaria-naïve subjects.^20^ To achieve a highly functional vaccine, antibodies targeting Pfs230 must be potent enough to block transmission of parasites to mosquitoes. Thus, understanding whether and how formulation with different adjuvants shapes the human antibody repertoire in response to Pfs230D1 is important to define B cell immunity to TBVs. Antibody repertoire analyses by B cell sequencing have been useful to determine common signatures in antibody genes and gain insights into the adaptive immune response to other vaccines.^21^ Here, we investigated the Pfs230D1 antibody repertoire at the antigen-specific, single cell level, in response to Alhydrogel® or AS01 adjuvants (Pfs230D1-EPA/Alhydrogel® and Pfs230D1-EPA/AS01).

Adjuvants can alter the selection of TCR responses, activate human B cells through TLR ligands,^22^ and increase the number of antigen-specific sequences in the BCR repertoire.^23^ In a mouse model of malaria vaccination, antibody repertoire sequencing determined that a TLR agonist adjuvant increased antibody variability in the animals and improved not only parasite neutralization but also efficacy against heterologous *Plasmodium* strains.^24^ AS01 adjuvant can promote robust immune responses during vaccination with CSP, the target antigen in the recently *P. falciparum* pre-erythrocytic vaccine RTS,S; vaccination with CSP virus-like particles formulated in AS01 (RTS,S/AS01E vaccine) induces high anti-CSP serum levels in malaria naïve subjects^25^ and confers protection against clinical malaria in young African children.^26^

Recent data have shown that a delayed, fractional 3^rd^ dose of RTS,S/AS01 can increase protection against controlled malaria infection from 62.7% (0-1-2 month regimen) to 87.5% (0-1-7 month regimen).^27^ In a study of over 100 human monoclonal antibodies (mAbs) obtained from plasmablasts of subjects receiving full or fractional RTS,S/AS01 regimens, IGHV3 heavy chain genes were the most frequent in the response to RTS,S/AS01, especially IGHV3-30 and IGHV3-33.^28^ Furthermore, IGHV3 genes have been identified in functional mAbs isolated from repeated immunization of malaria-naïve volunteers with infectious *P. falciparum* sporozoites (PfSPZ Challenge) under chloroquine prophylaxis (PfSPZ-CVac).^29^ While our results showed that most of the functional AS01 mAbs belonged to IGHV3, any correlation between the IGHV3 and functional activity for Pfs230D1-EPA/AS01 or RTS,S/AS01 vaccines remains to be investigated, including any adjuvant-specific effect. IGHV3 genes were also predominant in response to intravenous inoculation of *P. falciparum* sporozoites with no adjuvant formulation^30,31^ or after naturally acquired infection.^32^

Although IGHV1-69 antibodies uniformly bound to recombinant Pfs230D1 and showed high rates of IGHG mutations, none of them blocked parasite transmission when tested by mosquito in vivo assays. Although V gene usage by itself cannot predict whether an antibody will bind or not, IGHV1-69 genes are highly associated with cross-reactive binding in response to influenza vaccine.^33,34^ Another study showed that the IGHV1-69 gene is three times more abundant in mutated clusters compared to unmutated clusters. In fact, IGHV1-69 is one of the most polymorphic genes within the human IGHV gene cluster.^35,36^ IGHV1-69 polymorphisms seem to be substantially increased in African compared to Asian populations, for example,^37^ and our study was conducted among Malian subjects in west Africa. Curiously, IGHV1-69 was used in the sequence of LMIV230-01, a functional mAb isolated from a Malian adult enrolled in the same Alhydrogel trial as the subjects from the present study.^15^ Thus, we cannot exclude the fact that IGHV1-69 might be increased in clonally expanded cells and eventually generate functional sequences.

In our study CDR3 length was significantly longer in Alhydrogel mAbs sequences that bound to recombinant Pfs230D1 compared to those that did not bind. CDR3 length, however, was not associated with functional activity for any of the groups tested. Although we have identified a higher number of mutations in IGHG sequences in response to both vaccines [data not shown], a recent study has demonstrated that the number of mutations in heavy chain genes does not correlate with CDR3 length.^38^ In that study, antibody repertoire analyses in response to COVID-19 vaccines revealed that the average number of nucleotide mutations in IGH was 4.2 after the first dose and 11.7 after to the second dose, while the CDR3 length of IGH was unaltered after both doses.^38^

Our study has limitations. Time points of collection differed: Alhydrogel®, 14 days after dose 4; AS01, 7 days after dose 3. This was unavoidable for the AS01 samples because at the time of this study the samples after the 4^th^ vaccine dose were not yet available. However, this limitation was addressed in part by selecting samples with comparable antibody titers and serum functional activity between groups from the different trials (Supplementary Figure 3).

This work provides novel information on the diversity of antibody clonotypes elicited by vaccination with malaria TBVs and potential relationships between use of adjuvants and V gene usage, CDR3 length, binding to recombinant and native protein, and functional activity. Further research is required to explain why highly mutated IGHV1-69 sequences bind to recombinant Pfs230 but cannot access the epitope on the surface of the parasites.

## METHODS

### Human ethics statement

The clinical trials were approved by the FDA, by the ethics review boards from the Faculté de Médecine de Pharmacie et d’OdontoStomatologie (FMPOS), Bamako, Mali, and the US National Institute of Allergy and Infectious Diseases (NIH, Bethesda, MD, USA), as well as the Mali national regulatory authority. Written informed consent was obtained from study participants. The clinical trials are registered in clinicaltrials.gov (NCT02334462 for Pfs230D1-EPA/Alhydrogel and NCT02942277 for Pfs230D1-EPA/AS01) and the phase I in healthy adults was published elsewhere.^20^

### Human immunization and samples collection

Malian adults were vaccinated with four doses of 40 μg of Pfs230D1-EPA/Alhydrogel® or Pfs230D1-EPA/AS01 at planned study days 0, 28, 168, and 540. From the Pfs230D1-EPA/Alhydrogel® trial, sera and peripheral blood mononuclear cells (PBMCs) were obtained from eight participants, 14 days after the 4th dose (day 554). From the Pfs230D1-EPA/AS01 trial, collections occurred 7 days after the 3^rd^ dose. PBMCs (5 million cells per sample on average) were prepared for isolation of Pfs230D1-specific single memory B cells.

## ELISA

ELISA to assess total anti-Pfs230D1 IgG levels in sera from immunized subjects and evaluation of binding activity of human mAbs was performed as previously reported.^15^

### Identification and sorting of antigen-specific single memory B cells

Identification and sorting of Pfs230D1M-specific B cells was performed as previously described.^15^ Briefly, recombinant Pfs230D1M were chemically biotinylated using EZ-Link Sulfo-NHS-LC-Biotin (Thermo Fisher Scientific, Waltham, USA). The biotinylated proteins were then tetramerized with streptavidin labelled with PE (Prozyme, Hayward, USA). PBMCs from vaccinees were thawed in 37°C water bath, resuspended in complete Roswell Park Memorial Institute (RPMI) 1640 Medium with L-glutamine and 25 mM HEPES (Corning, Corning, NY, USA, 10-041-CV) and washed with phosphate-buffer solution (PBS) (Thermo Fisher Scientific, Waltham, MA, USA, 10010023). One μM of Pfs230D1 tetramers were added to the cells that were then incubated at 4ºC for 20 minutes. Cells were washed with PBS containing 10% fetal bovine serum (FBS) and incubated with 25 μL of anti-PE magnetic beads (Miltenyi Biotech, Bergisch Gladbach, Germany) for 25 minutes. Four mL of PBS were added to the solution, which was then passed over magnetized LS columns for elution of cell suspension enriched for antigen-specific cells. After enrichment with tetramer, PBMCs were stained with the following surface-conjugated antibodies: CD3 (UCHT1), CD14 (M5E2), CD56 (HCD56) Alexa Fluor 700, CD19 APC-CY7 (HIB19) CD20 PE-CY7 (2H7) purchased from Biolegend (San Diego, USA). The following gating strategy was used: singlet cells were selected for exclusion of non-B cells using CD3, CD14 and CD56 markers. Lymphocytes were gated for CD19+CD20+. Pfs230D1M-specific B cells were gated using PE and excluding non-Pfs230D1M cells gated using CF594, the fluorochrome used in the decoy BSA tetramer. Sorting was performed into 96 well plates using a FACSAria™ II instrument (BD Biosciences, San Jose, USA) with blue, red, and violet lasers. Pfs230D1M-specific memory B cells were analyzed according to the fluorescence staining profile described above and sorted directly into 96-well PCR plates using a 100 μM nozzle. After sorting, plates were immediately centrifuged at 1,278 x g for 30 seconds, transported in dry ice and stored at -80°C.

### Sequencing and data processing

Amplification of heavy and light chains and sequencing was performed by iRepertoire Inc. (Huntsville, AL, USA) as previously reported.^15^

### Standard Membrane Feeding Assay to assess functional activity of polyclonal antibodies in serum of vaccinees

Transmission Reduction Activity (TRA) was assessed by the estimating the reduction of *P. falciparum* burden in infected-mosquitoes fed with intact sera from vaccinees, using Standard Membrane Feeding Assay (SMFA), as previously reported.^20^

### BCR Sequence Analyses

2,393 sequences were assigned Ig genes, out of 2,397 input sequences (99.83%). Gene assignment and other downstream analysis were performed with the Immcantation suite container version 4.1.0, which includes IMGT reference germlines downloaded on 2020-08-12, IgBLAST version 1.16.0, Change-O 1.0.0, Alakazam 1.0.2 and SHazaM 1.0.2. After gene assignment with IgBLAST, 126 unproductive sequences were removed. Data were processed to retain one heavy chain sequence and at most two light chain sequences per well. In wells with multiple heavy chain sequences, we retained the one with the highest cdr3_read_count, and required it to account for more than 50% of the total heavy chain cdr3_read_count in the well. In wells with more than two light chain sequences, we retained the two sequences with the highest cdr3_read_count, each accounting for more than 25% of the total light chain cdr3_read_count in the well, and together for more that 75% of the total light chain cdr3_read_count in the well. Sequences from wells presenting light chain only were removed from the analysis. One well was excluded for possible contamination (subject 2, well A07, Pfs230 Alhydrogel). A total of 1,345 sequences passed these filters (733 heavy, 612 light), and constitute the final set of sequences analyzed, with an average of 57 sequences per subject for the Alhydrogel group, and 60 sequences for the AS01 group. Most of the sequences (79.26%) from the heavy chain sequences correspond to VH:VL pairs (Supplementary Figure 4).

### Sequence selection, mAbs expression and purification

VH and VL sequences of Pfs230D1-specific single memory B cells were selected for expression based on evidence of clonal expansion. If multiple cells contained identical CDR3 sequences, it was attributed to clonal expansion, and thus evidence of activation. Naturally paired VH/VL sequences were expressed in an IgG1 backbone by LakePharma Inc. Human mAbs were expressed using 0.1 L transient production in HEK293 cells (Thermo Fisher Scientific) and purified using protein A affinity chromatography. Purified proteins were submitted to sterile filtering using a 0.2 μm sterile filter and characterized for more than 90% purity by capillary electrophoresis-SDS, concentration, and endotoxin according to the manufacturer’s procedures.

### Standard Membrane Feeding Assay (SMFA) to test functional activity of mAbs

SMFA was performed to assess the ability of mAbs to block the development of *P. falciparum* strain NF54 oocysts in the mosquito midgut, as previously reported.^39^

All the mAbs from the AS01 group were tested at 100 μg/mL. The mAbs from the Alhydrogel group were initially screened at 375 μg/mL, and then the mAbs with >65 % TRA at 375 μg/mL were further tested at 100 μg/mL. A functional mAb was defined as a mAb with > 75% TRA at 100 μg/mL.

### Statistical analyses

Statistical analyses of AIRR-seq were performed using Wilcoxon rank sum test. The Benjamini–Hochberg false discovery rate (FDR) was used to correct for multiple hypothesis testing and the criteria for significance was p<0.05 and FDR <0.20. The descriptions of the tests used are contained within the figure legends.

## AUTHOR CONTRIBUTIONS

CHC and PED conceived the study. SH, IS, and PED conceived the clinical protocols and supervised the clinical trials. CHC, BCNT, AB, and KM performed experiments. CHC, SM, AB, and BCNT performed data analyses. CL supervised the SMFA experiments. SHK supervised the Bioinformatic analyses. CHC wrote the manuscript. All authors interpreted the data and revised the manuscript.

## ACKNOWLEDGMENTS

This work was funded by the Intramural Research Program of the National Institute of Allergy and Infectious Diseases, National Institutes of Health and partially funded by PATH (grant 01665623-GEN from PATH’s Malaria Vaccine Initiative to PED for BCR analysis and hmAb production). We are thankful to the study participants and the Clinical and Immunology teams in the Laboratory of Malaria Immunology and Vaccinology at NIAID. J. Patrick Gorres proofread the manuscript. Olga Muratova and Edward Owen provided assistance on parasite culture. This study was conducted in collaboration with GlaxoSmithKline Biologicals SA, with coordination and scientific input by Marc Lievens and Danielle Morelle during the design and execution of the NCT02942277 clinical trial. GlaxoSmithKline Biologicals SA was provided the opportunity to review a preliminary version of this manuscript for factual accuracy, but the authors are solely responsible for final content and interpretation.

## COMPETING INTERESTS

CC is a consultant for USAID. SHK receives consulting fees from Northrop Grumman and Peraton.

## Notes

### Competing Interest Statement

CHC is a consultant for USAID. SHK receives consulting fees from Northrop Grumman and Peraton

