## Supplementary material for "Antibody gene features associated with binding and functional activity in vaccine-derived human mAbs targeting malaria parasites": Sup. material

**Contents:**

Supplementary Figures 1-4
Supplementary Table 1

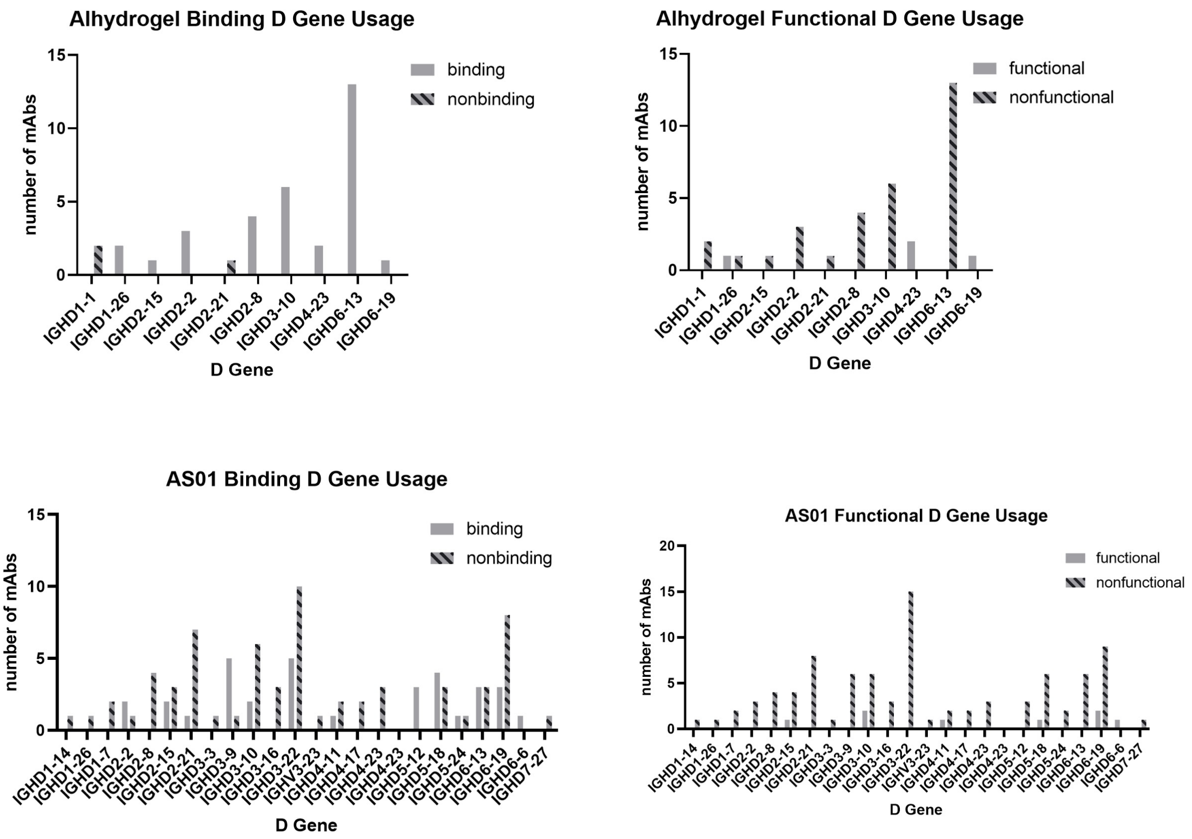


**Supplementary Figure 1– D genes present in mAbs, grouped by binding and functional activity profile.** Binding was assessed by ELISA and functional activity by SMFA. Functional antibodies are reported as with functional activity higher than 75% at 100µg/mL.

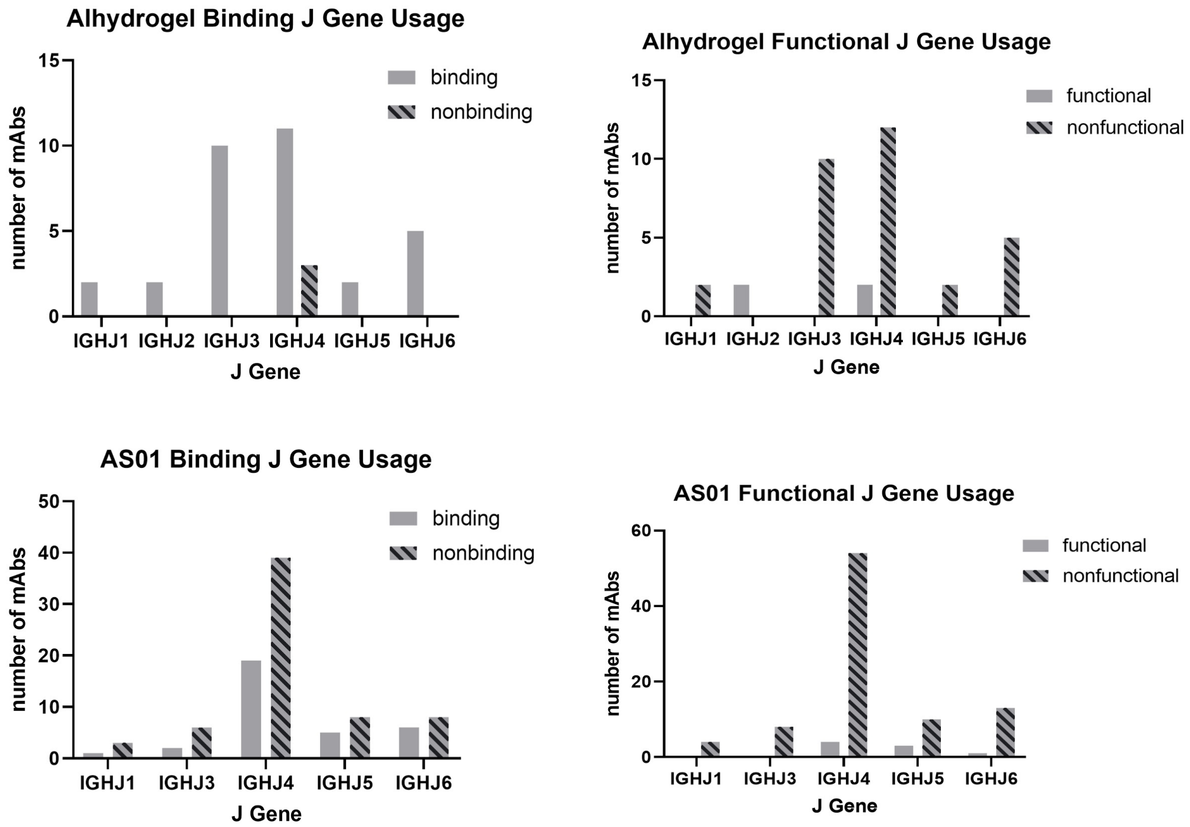


**Supplementary Figure 2 – Heavy chain J genes present in mAbs, grouped by binding and functional activity profile.** Binding was assessed by ELISA and functional activity by SMFA. Functional antibodies are reported as with functional activity higher than 75% at 100µg/mL.

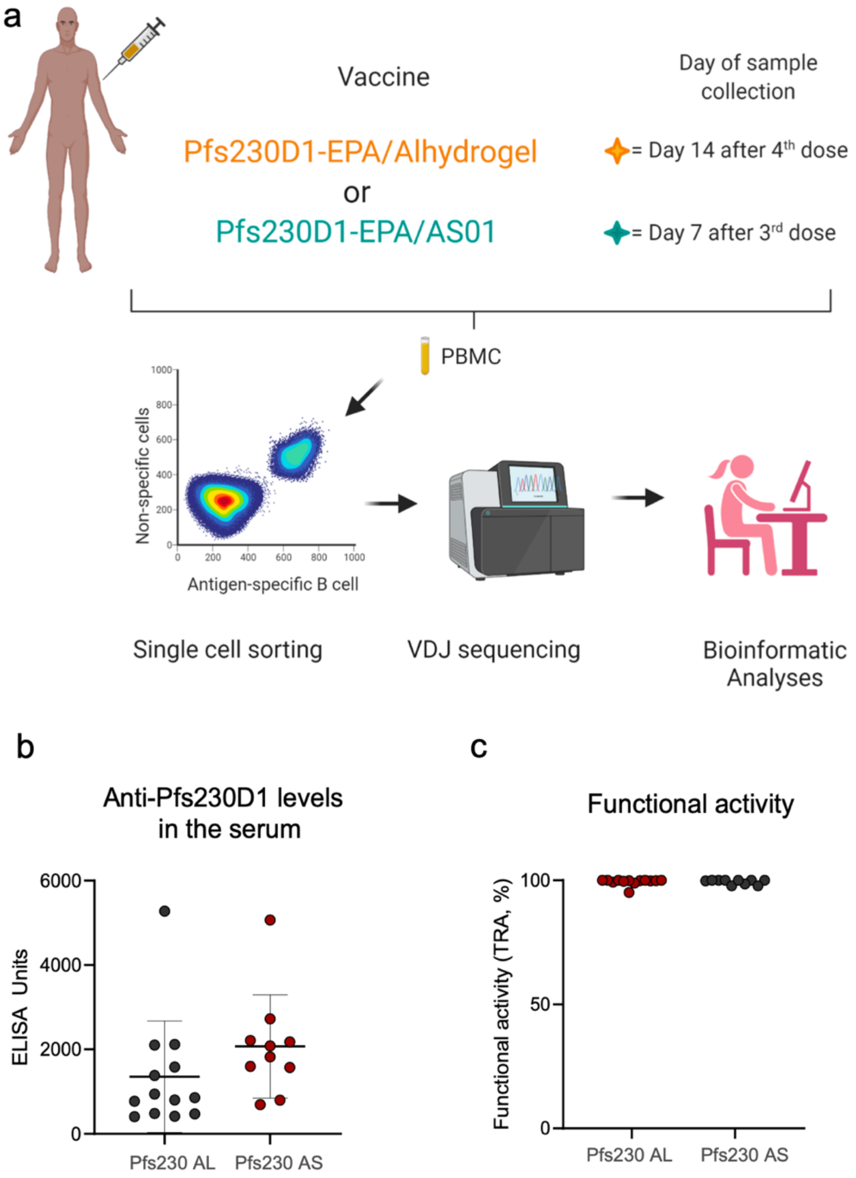


**Supplementary Figure 3- Antibody repertoire in Pfs230D1-specific single B cells in response to vaccination with Alhydrogel or AS01 adjuvants. (a)** Malian adults received 3 or 4 doses of Pfs230 conjugated with the carrier Exoprotein A and formulated with either Alhydrogel® or AS01 adjuvants. PMBCs were collected from subjects receiving Pfs230D1-EPA/Alhydrogel® (Pfs230AL) or Pfs230D1-EPA/AS01 (Pfs230 AS) and Pfs230D1-specific single B cells were sorted and had their B cell receptor sequenced. Bioinformatic analyses were performed using the Immcantation framework. Samples from the subjects enrolled in the clinical trial with vaccines formulated with Alhydrogel® were collected 14 days after the 4$^{th}$ dose, and with AS01, were obtained 7 days after the 3$^{rd}$ dose **(b)** Anti-Pfs230D1 IgG titers in response to both vaccines were measured by ELISA. **(c)** Serum functional activity was assessed by SMFA and determined by the ability to reduce the number of oocysts in midguts of infected *Anopheles* mosquitoes fed with *NF54 Plasmodium falciparum*.

a

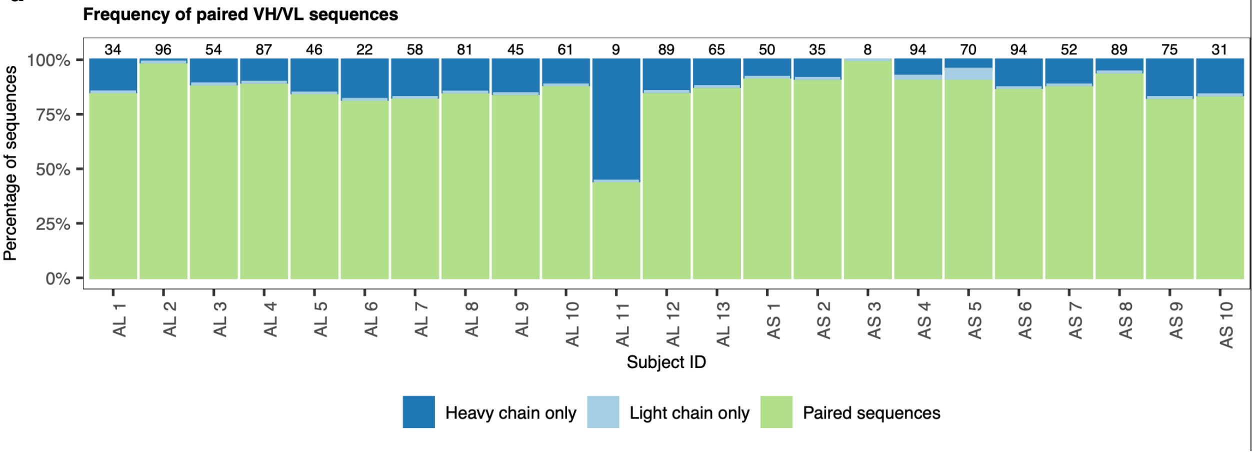


**Supplementary Figure 4 -** Proportion of Heavy chain (VH), light chain (VL) and pairs of VH/VL. Values in the Y axis represent the percentage for each of these three groups of sequences.

### SUPPLEMENTARY MATERIAL (TABLES)

| Alhydrogel (Subject ID) | Initial number sequences | Number of sequences after filtering | AS01 (Subject ID) | Initial number of sequences | Number of sequences after filtering |
|---|---|---|---|---|---|
| **1** | *69* | *34* | **1** | *69* | *50* |
| **2** | *183* | *96* | **2** | *55* | *35* |
| **3** | *112* | *54* | **3** | *16* | *8* |
| **4** | *148* | *87* | **4** | *152* | *94* |
| **5** | *98* | *46* | **5** | *123* | *70* |
| **6** | *71* | *22* | **6** | *119* | *94* |
| **7** | *106* | *58* | **7** | *78* | *52* |
| **8** | *154* | *81* | **8** | *165* | *89* |
| **9** | *82* | *45* | **9** | *109* | *75* |
| **10** | *125* | *61* | **10** | *52* | *31* |
| **11** | *37* | *9* | | | |
| **12** | *157* | *89* | | | |
| **13** | *117* | *65* | | | |
| TOTAL: | ***1459*** | ***747*** | TOTAL: | ***938*** | ***598*** |

**Supplementary Table 1 – Number of total BCR sequences per subject, including both VH and VL sequences.**
